# Deleterious Mutations Accumulate Faster in Allopolyploid than Diploid Cotton (*Gossypium*) and Unequally Between Subgenomes

**DOI:** 10.1101/2021.09.22.461419

**Authors:** Justin L. Conover, Jonathan F. Wendel

## Abstract

Whole genome duplication (polyploidization) is among the most dramatic mutational processes in nature, so understanding how natural selection differs in polyploids relative to diploids is an important goal. Population genetics theory predicts that recessive deleterious mutations accumulate faster in allopolyploids than diploids due to the masking effect of redundant gene copies, but this prediction is hitherto unconfirmed. Here, we use the cotton genus (*Gossypium*), which contains seven allopolyploids derived from a single polyploidization event 1-2 million years ago, to investigate deleterious mutation accumulation. We use two methods of identifying deleterious mutations at the nucleotide and amino acid level, along with whole-genome resequencing of 43 individuals spanning six allopolyploid species and their two diploid progenitors, to demonstrate that deleterious mutations accumulate faster in allopolyploids than in their diploid progenitors. We find that, unlike what would be expected under models of demographic changes alone, strongly deleterious mutations show the biggest difference between ploidy levels, and this effect diminishes for moderately and mildly deleterious mutations. We further show that the proportion of nonsynonymous mutations that are deleterious differs between the two co-resident subgenomes in the allopolyploids, suggesting that homoeologous masking acts unequally between subgenomes. Our results provide a genome-wide perspective on classic notions of the significance of gene duplication that likely are broadly applicable to allopolyploids, with implications for our understanding of the evolutionary fate of deleterious mutations. Finally, we note that some measures of selection (e.g. dN/dS, **π**_N_/**π**_S_) may be biased when species of different ploidy levels are compared.

## Introduction

Genome duplication (polyploidy) is among the most dramatic mutational processes in nature, causing myriad saltational changes at the cellular and organismal levels (Doyle and Coate 2019; Bomblies 2020; Fernandes Gyorfy et al. 2021), and is associated with consequential phenomena ranging from crop domestication (Renny-Byfield and Wendel 2014; Qi et al. 2021) to cancer progression (Matsumoto et al. 2021). Polyploidy is especially common in the angiosperms, with all extant species having experienced at least one or more polyploidy events during their evolutionary history (Jiao et al. 2011), and at least 30% of currently recognized species having a polyploidy event in the recent past (One Thousand Plant Transcriptomes Initiative 2019).

Novel evolutionary patterns created by polyploidy at the genic (e.g. neofunctionalization, subfunctionalization, and loss (Kuzmin et al. 2021)) and genomic (e.g., homoeologous recombination (Mason and Wendel 2020)) levels have been well documented across taxa, including the frequent asymmetry of these responses with respect to co-resident genomes in a polyploid nucleus. Nonetheless, many questions remain regarding the effects of natural selection on polyploid relative to diploid genomes (Baduel et al. 2019; Monnahan et al. 2019) and the interplay between these novel evolutionary patterns and the long-term trajectories of genome evolution (Qi et al. 2021) following polyploidization (e.g. biased fractionation).

One of the earliest predictions about natural selection in polyploids relative to diploids is that putatively deleterious mutations may accumulate faster due to the masking effect of completely or partially recessive deleterious mutations in duplicated genes (Haldane 1932; Hill 1970; Bever and Felber 1992). Only recently, however, have these predictions begun to be evaluated in young allopolyploids such as *Arabidopsis kamchatica (Paape et al. 2018)* and *Capsella bursa-pastoris (Douglas et al. 2015; Kryvokhyzha, Salcedo, et al. 2019; Kryvokhyzha, Milesi, et al. 2019)*, and autotetraploid *Arabidopsis arenosa* (Monnahan et al. 2019). Because the number of deleterious mutations is strongly influenced by shifts in demography and mating system (Brandvain and Wright 2016), which may coincide with polyploid formation (Grant 1981; Barringer 2007), a clear link between ploidy level and the accumulation of deleterious mutations is challenging to demonstrate in natural polyploid populations.

The cotton genus (*Gossypium*) represents one of the best studied allopolyploid systems (Wendel and Grover 2015; Hu et al. 2021). The genus includes approximately 45 currently recognized diploid species classified into eight genome groups (A-G, and K), and seven allopolyploid species resulting from a single (Grover et al. 2012) allopolyploidization event ~1-2 million years ago between members of the A and D genome groups (Wendel 1989). Although the most closely related extant species of these two progenitor diploids are found in southern Africa and Northern Peru, respectively, the polyploids are only found in the Americas (Figure 1). Most wild populations, including those of the two independently domesticated species *G. hirsutum* (AD_1_) and *G. barbadense* (AD_2_), occur in small, isolated populations on islands or in coastal regions. Subsequent to their initial domestication 4,000 - 8,000 years ago in the Yucatan Peninsula (AD_1_) and NW S. America (AD_2_), respectively, the ranges of the two domesticated species rapidly expanded to encompass much of the American tropics and subtropics and then spread globally with the rise of the international cotton fiber trade (Yuan et al. 2021).

**Figure 1:**
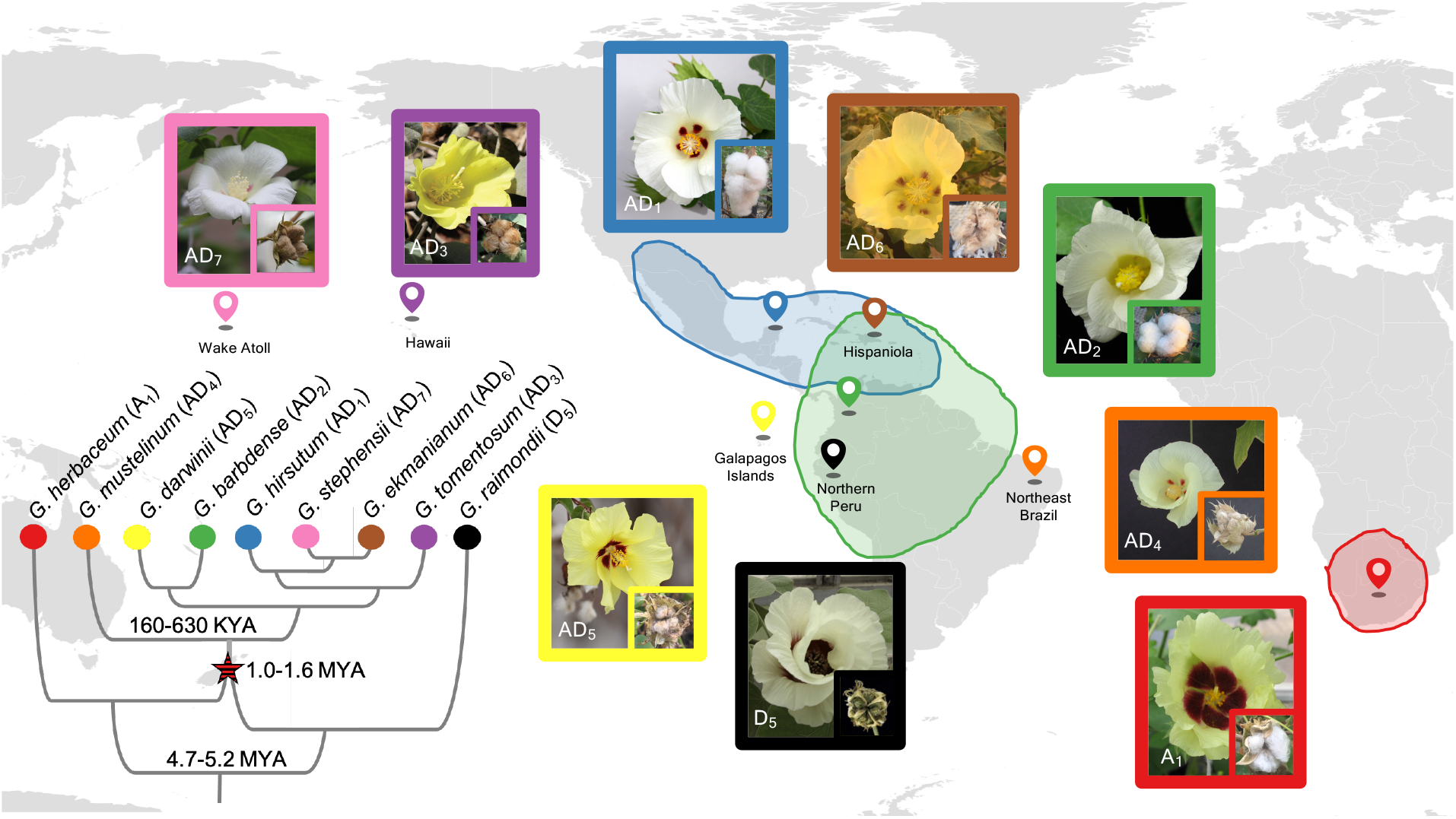
Phylogeny and Biogeography of *Gossypium* Allopolyploids and Progenitor Diploids. Diploid *Gossypium* species are classified into eight diploid genome groups. The A (represented by *G. herbaceum*) and D (represented by *G. raimondii*) genome groups diverged approximately 5 million years ago (MYA), with ranges in different hemispheres. Allopolyploids formed *circa* 1-1.6 MYA following transoceanic dispersal of an A genome ancestor (modeled by *G. herbaceum* (A_1_)) to the Americas and hybridization with a native D genome species (modeled by *G. raimondii* (D_5_)). Subsequent diversification of the new allopolyploid (AD genome) lineage led to the evolution of seven currently recognized species with a broad geographic range in the Americas and the Pacific islands. Flower and fruit morphology for each species is shown, and the island location and geographic range is indicated. Branch lengths on the phylogeny are not to scale but notable divergence times are labeled.

Here we describe the evolutionary trajectory of deleterious mutations in two wild diploid and six wild allopolyploid cotton species (all descended from a single allopolyploidization event), with a focus on how allopolyploidization and speciation have shaped the number and genomic distribution of deleterious mutations. We use two methods to estimate the strength of selection at the amino acid and nucleotide level and show support for a nearly century-old hypothesis that polyploids accumulate mutations faster than their diploid progenitors. We demonstrate that, in agreement with this hypothesis, polyploidy has the greatest influence on strongly, rather than moderately or mildly, deleterious mutations. We also find that deleterious mutations accumulate asymmetrically between the two co-resident subgenomes in the allopolyploid nucleus, indicating that these masking effects may act unequally between the subgenomes. In total, our results support theoretical predictions that allopolyploidy can lead to a faster rate of deleterious mutation accumulation through masking of recessive deleterious variants, and that the relationship of the rate of deleterious mutation accumulation between subgenomes and their progenitor diploids is complex, even when comparing identical pairs of single-copy homoeologs among lineages.

## Results

### Patterns of Synonymous and Nonsynonymous Mutations

To investigate patterns of deleterious mutations, we viewed our data at three phylogenetic depths: SNPs segregating within the global phylogenetic tree (Figure 2A-D), SNPs that emerged since the divergence of each subgenome from its respective diploid progenitor (Figure 2E-H), and SNPs that are still variable within the polyploids (Figure 2I-L). Each group is a subset of the previously described group. We restricted our analyses to a set of 8,884 single-copy, syntenically conserved homoeologous pairs of genes (17,768 genes in total) that showed no evidence of gene loss, gene copy variation, tandem duplication, ambiguous read mapping, homoeologous exchange, or homoeologous gene conversion (See Methods; Supplementary Figure 1). Notably, the patterns described below are largely reflected in genome-wide patterns as well (Supplementary Figure 2), indicating that filtering criteria did not bias overall results, and that, in cotton, homoeologous interactions have minimal effects on subgenome-specific SNP patterns (Supplementary Figure 1).

**Figure 2:**
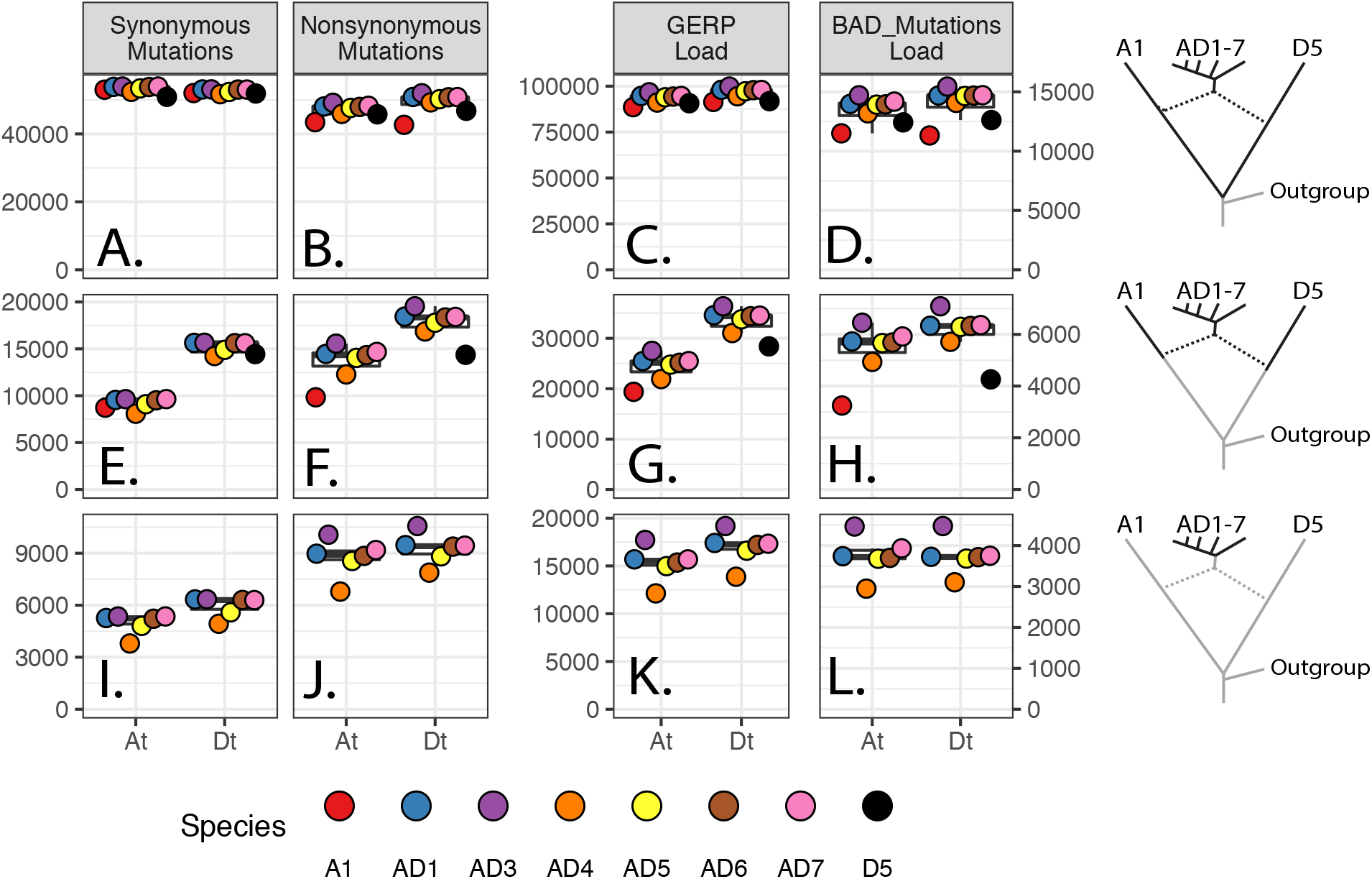
Derived Mutations and Deleterious Loads at Three Phylogenetic Depths. Number of derived synonymous, nonsynonymous, and deleterious mutations in the CDS regions of 8,884 pairs of homoeologs (17,768 genes in total) in eight cotton species at three phylogenetic depths (indicated by bold branches in phylogeny at right). For all panels, the ancestral state of each SNP was determined using three Australian cotton species as an outgroup (see Methods). The deepest phylogenetic depth **(ABCD)** includes all derived mutations that originated since the divergence of the A and D diploid progenitors; the middle row **(EFGH)** shows SNPs that are variable within each subgenome and its associated progenitor diploid species; and the bottom row **(IJKL)** shows SNPs that originated post-polyploidy and are variable within the polyploids. **(AEI)** Synonymous mutations. **(BFJ)** Nonsynonymous mutations. The y-axis for both synonymous and nonsynonymous is shown at left, and represents the sum of the derived allele frequencies, interpreted as the average number of derived SNPs in that category in each species. **(CGK)** GERP Load of each species, calculated as the sum of (derived allele frequency * GERP Score) for all SNP positions with GERP > 0. **(DHL)** Number of deleterious mutations in each species, calculated by BAD_Mutations with Bonferroni-corrected significance (see Methods). Y-axis represents the sum of the derived allele frequencies, and indicates the average number of deleterious mutations in each species at a given phylogenetic depth. **Note:** for **(EFGH)**, comparisons between subgenomes cannot be made because the D5 diploid is more distantly related to the D subgenome than the A1 diploid is related to the A subgenome. Therefore, we would expect a larger number of derived mutations in D than A simply due to evolutionary history rather than to polyploidization *per se*.

Using the curated set of 8,884 pairs of homoeologous genes, we found no evidence for differences in the rate of synonymous mutation accumulation in diploids versus polyploids at any phylogenetic depth (Figure 2A, 2E), although differences can be found within the polyploid species (Figure 2I), with *G. mustelinum* (AD_4_, Orange) and *G. darwinii* (AD_5_, Yellow) having consistently lower rates than the rest of the clade, and in both subgenomes. When viewing SNPs that have accumulated since the divergence of the earliest polyploid lineage (Figure 2AI), there is an asymmetry between subgenomes in the rate of synonymous site changes, with the Dt (“t” denoting “tetraploid”) subgenome containing a moderately higher number of synonymous mutations than the At subgenome for all species. This difference potentially indicates a higher mutation rate or relaxed background selection in genic regions of the Dt subgenome compared to homoeologous genic regions of the At sugenome, and is consistent with previous analysis finding that genes in the Dt subgenome are evolving faster than genes in the At subgenome in five allopolyploid cottons (Chen et al. 2020).

In contrast to the relative homogeneity in rates of synonymous substitution among diploids and polyploids, rates of nonsynonymous mutation accumulation differed significantly at all phylogenetic depths. Notably, at the deepest phylogenetic depth (Figure 2B), estimates for the number of derived nonsynonymous mutations in the diploids *G. herbaceum* (A_1_, Red) and *G. raimondii* (D_5_, Black) did not differ between subgenomes, indicating that any mapping biases or erroneous SNP calls in these samples were removed by our SNP filtering criteria. *Gossypium raimondii* (D_5_) contained more derived nonsynonymous mutations than did *G. herbaceum* (A_1_), and this lineage-specific difference was reflected in the Dt and At subgenomes as well. When lineage-specific effects that arose from the long, shared ancestry between the subgenomes and their progenitor diploids were removed (Figure 2F), a clear distinction between the rates of nonsynonymous mutations between diploids and their respective subgenomes in the allopolyploids becomes apparent. In all polyploids, the At subgenome contained between 25-58% more nonsynonymous mutations than *G. herbaceum* (A_1_, Red), and the Dt subgenome contained 17-36% more than *G. raimondii* (D_5_, Black). These results demonstrate that even though the rates of synonymous mutation accumulation did not differ significantly between the diploids and polyploids, polyploidy significantly increases the rate of nonsynonymous substitution accumulation. Finally, when restricting our attention to only those mutations that have arisen following polyploid formation (Figure 2J), the lineage-specific patterns observed for nonsynonymous mutations were largely identical to the patterns of synonymous mutations. For example, *G. mustelinum,* (AD_4_, Orange) consistently had the lowest number of derived mutations in both subgenomes. Notably, however, the Hawaiian Island endemic *G. tomentosum* (AD_3_, Purple) has a higher number of derived nonsynonymous mutations than expected based on the patterns of synonymous mutations, potentially reflecting the population bottleneck associated with its origin following long-distance dispersal to the Hawaiian Islands (see Discussion). In summary, polyploidy significantly enhances the rate of nonsynonymous mutation accumulation in all *Gossypium* allopolyploids, and does so asymmetrically across co-resident genomes.

### Polyploidy Increases Rate of Deleterious Mutation Accumulation

Because the fitness effects of most nonsynonymous mutations can vary widely, from neutral to lethal, we asked if the elevated rate of nonsynonymous mutations observed in polyploid *Gossypium* reflects an increase in neutral or nearly-neutral nonsynonymous mutations, or if instead this elevation is attributable to a greater accumulation of deleterious mutations. To address this, we used two approaches to estimate whether a mutation was deleterious: BAD_Mutations and GERP++. BAD_Mutations performs a likelihood ratio test from a gene-specific multi-species alignment to determine if a mutation at a particular nonsynonymous site is deleterious, while GERP++ uses a genome-wide multiple sequence alignment (i.e. agnostic to genic regions) to estimate the degree of conservation at a particular site in the genome (including noncoding and synonymous sites). Notably, because one of the hallmark long-term processes following polyploidy is pseudogenization (Wendel 2015), recently pseudogenized sequences may still display some degree of conservation across the multiple sequence alignment, but may not be inherently deleterious. Therefore, to avoid inflating estimates of deleterious mutations in polyploids compared to diploids, we used GERP only within the exonic regions of the 8,884 homoeologs. Additionally, while GERP can score the degree of deleteriousness of a mutation, BAD_Mutations can only classify variants into deleterious or not deleterious. Therefore, the values shown in Figure 2DHL represent the sum of the allele frequencies of derived deleterious mutations, similar to the values for Figure 2AEI and Figure 2BFJ. For analysis with GERP, we used the GERP load, which incorporates the deleteriousness of each variant into the score, summing the frequency of each derived allele multiplied by it’s GERP score (see (Rodgers-Melnick et al. 2015; Wang et al. 2017)).

As shown in Figure 2, both of the foregoing analyses demonstrate that deleterious mutations accumulate in polyploids in a manner similar to nonsynonymous mutations, suggesting that the difference in nonsynonymous sites cannot be wholly attributed to putatively neutral or nearly-neutral alleles. For example, there is remarkable consistency in the patterns of deleterious mutations that have accumulated since the divergence of the diploid from its respective diploid progenitor in both the count of nonsynonymous substitutions (Figure 2F), the GERP load (Figure 2G), and the number of deleterious mutations (Figure 2H). In all three columns, the diploids show fewer accumulated alleles than the polyploids, *G. tomentosum* (AD_3_, Purple) shows the highest number of all the polyploids, and *G. mustelinum* (AD_4_, Orange) shows the fewest of all the polyploids.

An interesting pattern arises when comparing estimates of the GERP load (Figure 2G) and number of deleterious mutations (Figure 2H) between diploids and their closely related subgenomes: while the total number of deleterious mutations in the At subgenomes was 52-99% higher in the polyploids than the diploids (Figure 2H), the GERP load in the polyploids was only 13-42% higher (Figure 2G). Similar patterns were found in the Dt subgenome, with 34-66% more deleterious mutations in the polyploids than the diploid, but only a 9-13% increase in GERP load. This discrepancy could reflect inherent differences in the types of sequences used and how deleteriousness is quantified between the two methods, suggesting that the use of multiple analytical tools for detection of genetic load may yield more nuanced insights than either method on its own (See Discussion).

### Asymmetries in the Rate of Deleterious Mutation Accumulation

Although deleterious mutations are accumulating faster in polyploids relative to diploids, it is not obvious whether this increased rate is different from the increased rate of accumulation of nonsynonymous mutations. To test this, we compared, among ploidy levels, the total proportion of nonsynonymous mutations that were considered deleterious by BAD_Mutations (Figure 3). For SNPs that originated since the divergence of the A and D diploids (Figure 3A), the proportion of nonsynonymous sites that are deleterious is roughly 2% higher in polyploids than in diploids, despite the shared evolutionary history of more than 4 million years between each subgenome and their respective diploid progenitors. Notably, as similarly shown in Figure 2, the proportion of nonsynonymous mutations that are inferred to be deleterious in both diploids is equivalent when mapped to either subgenome, indicating that our filtering criteria did not differentially exclude deleterious or non-deleterious SNPs with respect to which subgenome the diploid reads were mapped.

**Figure 3:**
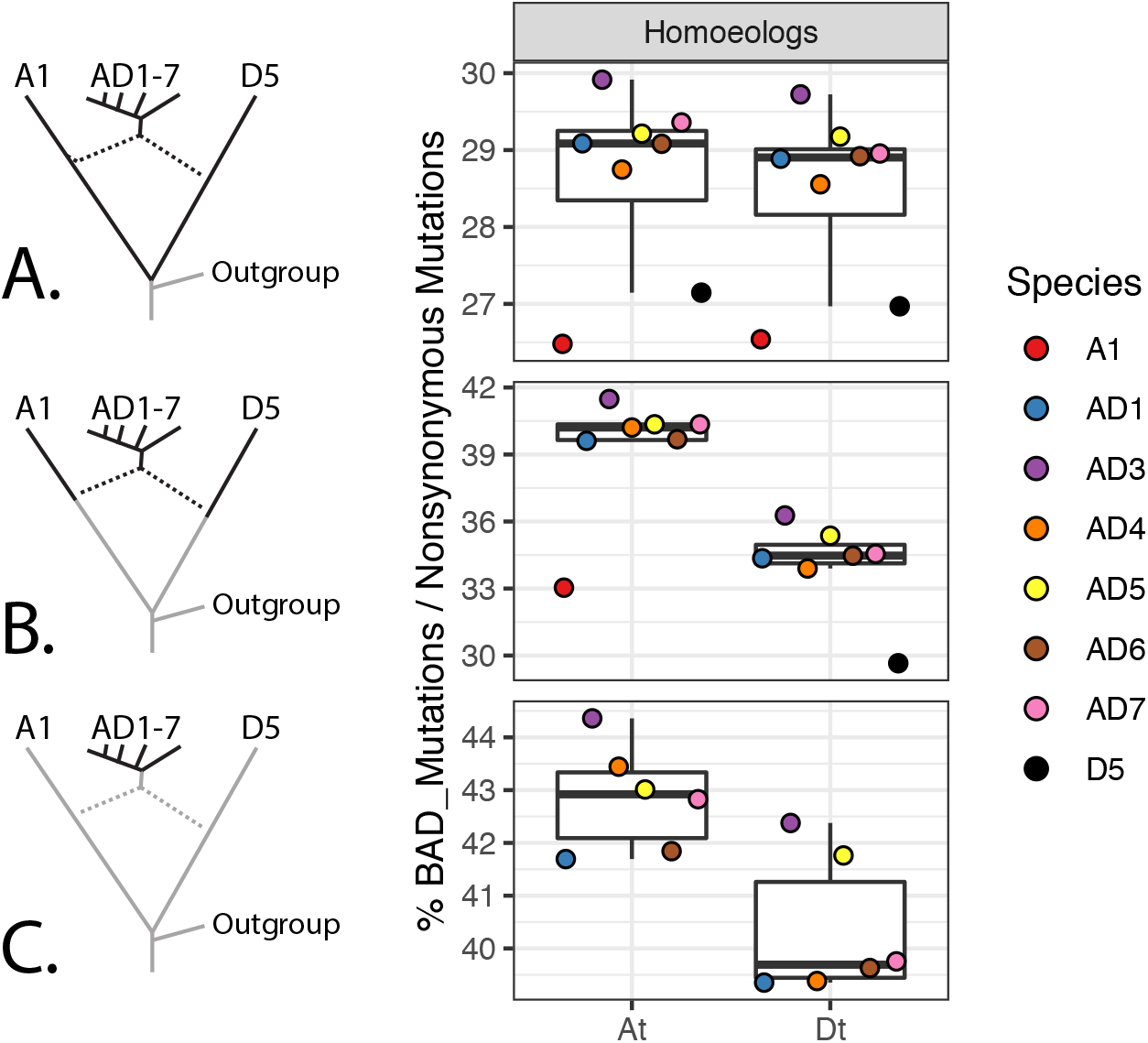
Proportions of All Nonsynonymous Mutations That Are Deleterious. Rows **A, B**, and **C** summarize SNPs segregating within the entire clade, within each subgenome and its respective progenitor diploid, and within each subgenome, as indicated by the bolded branches along the phylogeny at left. Values indicate the proportion of nonsynonymous SNPs that are deleterious within 8,884 homoeologous pairs (17,768 total genes) that are syntenically conserved between the two subgenomes of *G. hirsutum* (see Methods for filtering criteria). For example, the values in row A are calculated by dividing the values in Figure 2D by the values in Figure 2B for each species. **Note:** Similar to Figure 2, comparisons between subgenomes in row **B** reflect differing phylogenetic distances, not asymmetries between the subgenomes and/or their diploid progenitors.

At shallower phylogenetic depths (Figure 3B), the difference between diploids and polyploids becomes even clearer, with polyploids exhibiting 3-4% higher proportions of deleterious SNPs in the Dt subgenome and 5-12% higher in the At subgenome than their respective diploid progenitors. The most unbiased and straightforward comparison of the asymmetry in strength of purifying selection between the two subgenomes of allopolyploid cottons is provided by mutations that have occurred following polyploidization (Figure 3C). Here, the At subgenome of all species contain a 2-3% high proportion of deleterious SNPs than the Dt subgenome, indicating that differences exist in the strength of purifying selection between the two homoeologous subgenomes that have resided in the same nucleus for over a million years. This pattern is also observed when deleterious SNPs are mapped onto the phylogeny (Supplementary Figure 3). Additionally, there is more variation among species in the At subgenome than in the Dt subgenome, although the patterns in this respect are not simple. The amount of subgenomic asymmetry is smallest in *G. darwinii* (AD_5_, Yellow) from the Galapagos Islands, and largest in the Brazilian endemic and inland species *G. mustelinum* (AD_4_, Orange), indicating that asymmetries between subgenomes of the same species may vary within a single clade of allopolyploids.

### Disentangling Demography and Selection from Effects of Ploidy

Demography is a potential confounding factor in estimating the rate of deleterious mutation accumulation. Shifts in demography are known to complicate inferences of the strength of selection and genetic load (Brandvain and Wright 2016); for example, even in one of the best studied demographic shifts, the Out of Africa migration in humans, several papers (Lohmueller et al. 2008; Gazave et al. 2013; Simons et al. 2014; Henn et al. 2016; Simons and Sella 2016) have reached seemingly contradictory conclusions on whether genetic load has increased as a result of these shifts in demography (but see (Lohmueller 2014)). The pattern of deleterious mutation accumulation has also been well-documented in bottlenecks and population growth associated with domestication in crops such as maize (Wang et al. 2017), soybean and barley (Kono et al. 2016), sorghum (Lozano et al. 2021), cassava (Ramu et al. 2017), and rice (Liu et al. 2017).

Polyploidy is typically associated with a population bottleneck (Grant 1981; Barringer 2007), but because the genetic diversity of both the diploid and polyploid species in this study is low (Table 1), demographic modeling of the depth or duration of population bottlenecks and range expansion following polyploid formation is not straight-forward. Generalized patterns of the effects of demography on deleterious mutations, however, can serve as a null expectation to test if our data follows the same trends observed under varying demographic scenarios, as explained in the following.

**Table 1:** Nucleotide Diversity (π) in 8,884 Homoeologs in Eight *Gossypium* Species, By Subgenome.

| Species | Species Code | At Subgenome | Dt Subgenome |
| --- | --- | --- | --- |
| *G. herbaceum* | A1 | 7.41E-04 |  |
| *G. raimondii* | D5 |  | 2.36E-04 |
| *G. hirsutum* | AD1 | 6.69E-04 | 7.06E-04 |
| *G. tomentosum* | AD3 | 1.75E-04 | 1.67E-04 |
| *G. mustelinum* | AD4 | 2.64E-04 | 3.15E-04 |
| *G. darwinii* | AD5 | 1.71E-04 | 1.60E-04 |
| *G. ekmanianum* | AD6 | 7.75E-04 | 7.67E-04 |
| *G. stephensii* | AD7 | 4.94E-05 | 5.59E-05 |

Demographic shifts, including population bottlenecks and expansions, have a large influence on the accumulation of deleterious mutations. According to the nearly neutral theory (Ohta 1992), the fate of deleterious mutations is determined by genetic drift instead of selection when the selection coefficient (s) of deleterious mutations is less than or equal to 1/(2N_e_), where N_e_ is the effective population size. The reduction of N_e_ during a population bottleneck would therefore allow weakly deleterious mutations to escape purifying selection (i.e. to behave as if they were neutral), while strongly deleterious mutations with a selective coefficient greater than 1/(2N_e_) would still be removed by purifying selection. On the other hand, as N_e_ increases during population expansion, mutations that are mildly deleterious are expected to be more efficiently purged from the population.

In both demographic scenarios, we expect that mildly or moderately deleterious mutations would be most differentially affected, while strongly deleterious mutations would consistently be removed by purifying selection. Based on this theory, if the differences in the number of deleterious mutations we see between diploids and polyploids are due to demography, then we would expect to see most of that difference reflected in mildly, rather than strongly, deleterious mutations. In contrast, if masking of deleterious alleles in polyploids is driving a higher rate of accumulation relative to diploids, this pattern will not be observed.

To test if our data were consistent with changes in demography, we first asked if there was a correlation between the degree of deleteriousness of a mutation (as measured by GERP) and its relative increase in the polyploids compared to the diploids. To answer this question, we plotted the relative change of deleterious mutations in each subgenome relative to its most closely related diploid progenitor. We plotted this relative change for three different degrees of deleteriousness - strongly deleterious mutations (4 < GERP ≤ 6), moderately deleteriousness (2 < GERP ≤ 4), and mildly deleteriousness (0 < GERP ≤ 2) deleterious (Figure 4). We found that in both subgenomes of all six polyploids, when comparing SNPs that had originated after the divergence of the diploid from its respective subgenome in the allopolyploids, strongly deleterious mutations accumulated at a faster rate relative to diploids than did moderately or mildly deleterious mutations, which is inconsistent with expectations under a demographic change model alone. We also observed this change under both an additive and recessive model of dominance (Supplementary Figure 5). In total, the rate of accumulation among mutations with different inferred degrees of deleteriousness do not suggest that the patterns we see can be explained solely by demographic changes, but that the masking effect of duplicated genes may play an important role in the determining the fate of deleterious mutations in allopolyploids.

**Figure 4:**
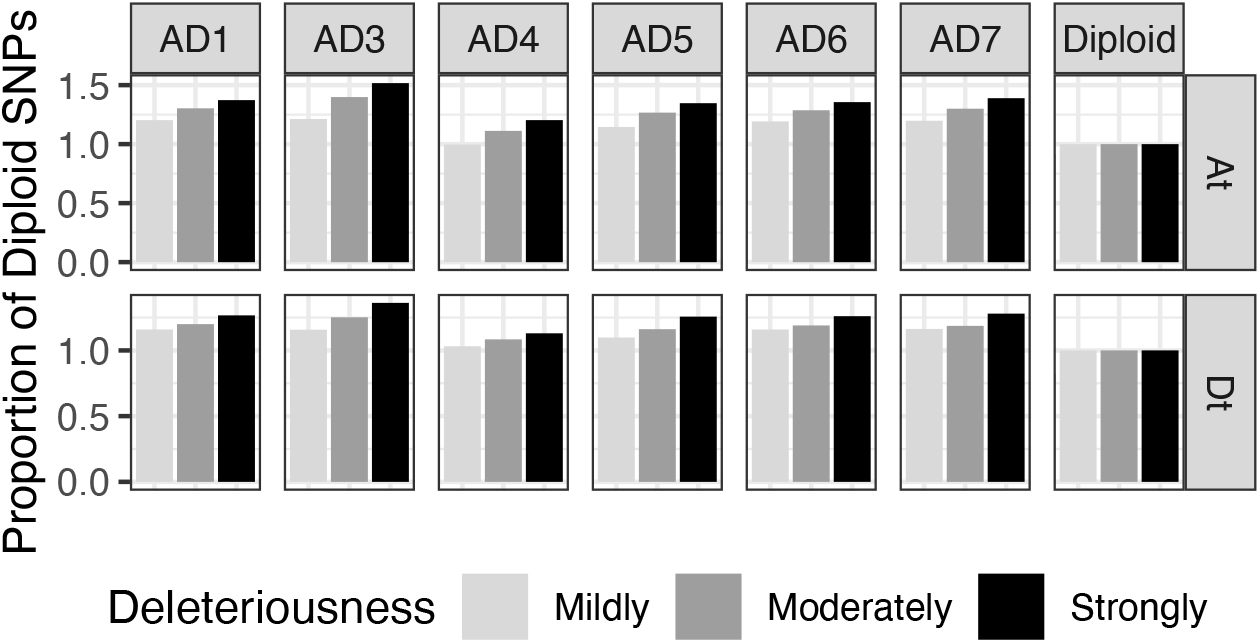
Relative Increase Of Deleterious Mutations Among GERP Categories in Polyploids Compared to Diploids. For SNPs that originated since the divergence of each subgenome from its diploid progenitor, we plotted the relative increase in deleterious alleles across three GERP load categories: mildly deleterious (0<GERP≤2; light gray), moderately deleterious (2<GERP≤4; gray), and strongly deleterious (4<GERP≤6; black). We used the diploid as the reference population, meaning that the relative increase of GERP load in the diploid is always equal to one for all categories. In both subgenomes of all polyploids, strongly deleterious mutations had the greatest relative increase compared to the diploids, followed by the moderately deleterious mutations, and finally, mildly deleterious mutations. This pattern does not fit the expected patterns under demographic models alone, where most of the changes between two populations should be seen in mildly or moderately deleterious mutations. However, under a model where recessive deleterious mutations are masked by their homoeologs, we would expect that strongly deleterious mutations would accumulate faster than moderately or mildly mutations (i.e the pattern we see here) due to the correlation between the recessivity of a mutation (h) and its selection coefficient (s).

## Discussion

### Effects of Polyploidy on Deleterious Mutation Accumulation

One of the earliest hypotheses regarding mutation accumulation in allopolyploids dates back to Haldane (Haldane 1932) where he posits that in allopolyploids, “one gene may be altered without disadvantage, provided its functions can be performed by a gene in one of the other sets of chromosomes.” Allopolyploids are therefore predicted be able to tolerate a higher mutational load than their diploid relatives, and putatively deleterious mutations may accumulate faster in polyploids than in their diploid relatives due to the masking effect of recessive or incompletely dominant deleterious alleles. Here, we demonstrate that these predictions are true in allopolyploid cottons. All polyploids in *Gossypium* harbor more mutations at phylogenetically conserved sites than do their closest diploid progenitors, as determined by two different methods of detecting deleterious mutations. We also find that the proportion of all nonsynonymous mutations that are inferred to be deleterious is higher in polyploids than in their diploid progenitors and that polyploidy has the greatest effect on strongly deleterious (and, inferentially, more recessive (Eyre-Walker and Keightley 2007; Huber et al. 2018)) mutations. Thus, using the power of comparative phylogenetics and genomics combined with analytical methods for detection of deleterious mutations, we demonstrate confirmation of a nearly century old hypothesis regarding natural selection in allopolyploid organisms.

### Demography Alone Cannot Explain Patterns of Deleterious Mutations in Polyploids

Estimating the strength of natural selection and genetic load is notoriously challenging (Lohmueller 2014) and is complicated by shifts in effective population size (including bottlenecks and expansions), mating systems, and effective recombination rates, among other life-history and demographic factors (Brandvain and Wright 2016). Here we illuminate an additional relevant consideration, i.e., whole genome duplication. Yet many of the considerations for populations that are not in demographic equilibrium also apply to *Gossypium*. Diversification in the cotton tribe (*Gossypieae*) has been characterized by numerous long-distance dispersal events (Grover et al. 2017), including the one from Africa to the Americas 1-2 MYA that led to the evolution of allopolyploid *Gossypium*. We note that in the Hawaiian Islands endemic *G. tomentosum*, the total number of synonymous substitutions is not significantly different from the rest of the polyploids, but the number of nonsynonymous and deleterious mutations is significantly increased, suggesting that the genetic bottleneck associated with island dispersal has elevated the number of deleterious mutations compared to the rest of the polyploids.

While demographic changes upon polyploid formation have been shown to change the number and frequency of deleterious mutations in other systems (Douglas et al. 2015; Paape et al. 2018; Baduel et al. 2019; Kryvokhyzha, Salcedo, et al. 2019; Kryvokhyzha, Milesi, et al. 2019), we show here that the patterns of mutation accumulation in *Gossypium* cannot be explained by demography alone, and that the data are more consistent with the nearly century-old hypothesis that recessive deleterious mutations can accumulate faster in allopolyploids due to the masking effect of duplicated genes and lack of recombination between subgenomes (Haldane 1932). Specifically, we show that strongly (and, hence, more recessive (Morton et al. 1956; Mukai et al. 1972; Eyre-Walker and Keightley 2007; Agrawal and Whitlock 2011; Huber et al. 2018)) deleterious mutations accumulate faster in polyploids compared to diploids than moderately or mildly deleterious mutations, and that this pattern is inconsistent with demographic shifts or long-term change in population size (Figure 4, Supplementary Figure 5).

### Asymmetry in Subgenomes in the Distribution of Deleterious Mutations

One of the elegant attributes of a clade of allopolyploid genomes derived from a single polyploidization event is that they offer a remarkable natural experiment for comparing subgenomes that have resided within the same nucleus for, in the case of *Gossypium*, approximately 1.5 million years. Once an allopolyploid is established, each subgenome is subjected to identical external or population-level factors, including demography, mating systems, and environmental and ecological conditions, as well as internal cellular processes, including identical DNA replication and recombination machinery. These features remove many of the confounding factors that may influence the genetic load and provide a simple comparative context for revealing evolutionary forces that might differentially affect co-resident genomes or homoeologs.

An unexpected finding of our analyses is the striking asymmetry in the proportion of all nonsynonymous mutations that are inferred to be deleterious between the two subgenomes of all allopolyploid species in *Gossypium*. We found that the At subgenome of all species contains 2-3% more nonsynonymous mutations that are inferred to be deleterious (Figure 3) even when only considering mutations that have arisen following the earliest allopolyploid diversification events, and correcting for removing the biases of unequal phylogenetic distances to each subgenome’s model progenitor diploid. Our work adds to a growing recognition that the two co-resident subgenomes in cotton allopolyploids may be shaped asymmetrically by evolutionary processes, including interspecific introgression and selection under domestication (Fang, Wang, et al. 2017; Fang, Guan, et al. 2017; Chen et al. 2020; Yuan et al. 2021), and that this phenomenon also extends to other important allopolyploid crop plants, including wheat (Pont and Salse 2017; Jiao et al. 2018) and *Brassica (Tong et al. 2020)*.

Teasing apart the genesis of differential subgenomic responses to selection is rendered challenging by several factors independent of phylogeny. We note, for example, the relevant example of the recently formed allopolyploid *Capsella bursa-pastoris* and its diploid progenitors, where consistent asymmetries in genetic load are reported between the subgenomes (Kryvokhyzha, Salcedo, et al. 2019; Kryvokhyzha, Milesi, et al. 2019) the differences likely reflect the dramatically different mating systems of the progenitors, in which the subgenome with the higher genetic load originated from an obligate outcrosser, *C. grandiflora* (Ne = 800,000), whereas the subgenome with the lower genetic load derives from the predominantly selfing *C. orientalis* (Ne = 5000)(Douglas et al. 2015). In another recently formed (20-250 thousand years ago) allopolyploid, *Arabidopsis kamchatica*, no asymmetry in the distribution of fitness effects between subgenomes was found, although it was observed that each subgenome of the allopolyploid contained more neutral and fewer deleterious alleles than either of the diploid progenitors (Paape et al. 2018). It is unclear, however, whether this shift was due to allopolyploidy *per se* or if it reflects the transition from an obligate outcrossing to a mating system with some degree of inbreeding, with a concomitant purging of partially or completely recessive deleterious alleles, as shown in several other systems (Arunkumar et al. 2015; Roessler et al. 2019). In *Gossypium*, all species have similar mating systems and a canonical outcrossing floral morphology including highly exserted styles and stigmas. Population sizes often are small, however, likely leading to relatively high levels of generalized inbreeding. At present, however, no data exist that address these considerations.

### Polyploidy, Redundancy, and Fitness Effects

One possible interpretation of our results is that *Gossypium* polyploids are less fit than their closely related diploid progenitors because they harbor more deleterious mutations in their genomes, especially mutations that have already been driven to fixation. We note that an additional possibility is that mutations in polyploids that occur at phylogenetically conserved sites may not actually have a deleterious effect on fitness as they do in diploids. Inferring the genetic load of a population simply by counting the number of deleterious variants assumes that all alleles contribute independently to the total genetic load of a population. However, because of the functional overlap of duplicated genes and, in most cases, absence of recombination between homoeologous chromosomes in an allopolyploid, a recessive deleterious mutation can never be present in all four copies of a gene and thus may be invisible to selection because of the masking effect of its homoeologous partner.

An important takeaway from this study is that recessive deleterious mutations in allopolyploids, at least at some loci, may actually accumulate in a manner more similar to neutral mutations, presumably because of the lack of recombination between subgenomes and, hence, the inability of purifying selection to “see” the negative effects of these mutations. Because these recessive deleterious mutations escape the effects of purifying selection, many traditional tests for detecting selection (e.g. dN/dS, **π**_N_/**π**_S_) may be biased when comparing a polyploid to diploid because the polyploid would be expected to accumulate putatively deleterious sites more quickly (and maintain a higher genetic diversity at nonsynonymous sites) than their diploid relatives.

Another important implication of this finding is that allopolyploidy (or gene duplication in general) may play an important and underrecognized role in determining how selection acts on new mutations, notwithstanding the burgeoning literature on fates of duplicate gene evolution (Conant et al. 2014; Shi et al. 2020; Veitia and Birchler 2021). The evolutionary trajectory of new mutations will largely be dependent on the selection coefficient (s) acting on that locus, and the dominance coefficient (h), defined as the proportion of the fitness cost that a mutation harbors when in a heterozygous state. In allopolyploids, however, the evolutionary fate of new mutations may be determined not only by allelic dominance at that locus, but also by the interaction arising from the coexistence of its homoeologous locus, a term we call “homoeologous epistatic dominance”. The relationships between this homoeologous epistatic dominance, allelic dominance, and the selection coefficient are likely complicated and potentially heavily influenced by other biological considerations such as biased expression of homoeologs, sub- or neofunctionalization, and homoeologous recombination, among others. Moreover, notwithstanding these polyploidy-specific effects, even the genome-wide relationships between two of these factors, allelic dominance and the selection coefficient, have only been modeled using genomic data in the past few years (Huber et al. 2018).

Nonetheless, understanding how this homoeologous epistatic dominance impacts the fitness effects of new mutations is an unexplored aspect of polyploid genome evolution, and it is not yet clear whether this will equally affect advantageous and deleterious variants. How homoeologous epistatic dominance operates with respect to functional properties arising from considerations such as gene balance (Veitia and Birchler 2021), dosage effects (Conant et al. 2014), structural and functional entanglement (Kuzmin et al. 2020; Kuzmin et al. 2021), and inter-subgenomic *cis- and trans-* effects (Bottani et al. 2018; Hu and Wendel 2019) would seem to represent important avenues for understanding how natural selection operates differently in polyploids compared to diploids. From an applied perspective, these insights could be important in agriculture, particularly because so many of our most important crop plants have a recent history that includes polyploidy (Renny-Byfield and Wendel 2014), and segregating patterns of genome fractionation have the potential to serve as targets of selection in crop improvement (Hufford et al. 2021).

## Materials and Methods

### Plant Materials and Sequencing

We used whole genome sequencing data from 46 individuals in *Gossypium*, including between two and ten individuals from each of eight species. Included in our sampling was six polyploid species originating from a single polyploidization event 1-2 million years ago (Wendel 1989; Hu et al. 2021), two diploid species representing models of the genome donors to the allopolyploids (A and D), and three species from Australia that served as outgroups for polarizing SNPs into ancestral and derived states. These sequences were previously described (Yuan et al. 2021), and SRA codes for all 46 resequenced individuals are listed in Supplemental Table 1. For *G. hirsutum*, we randomly chose ten accessions that were classified in the “Wild” population from Yuan et al. (Yuan et al. 2021), and for the other species, we chose all accessions available that did not show evidence of being mislabeled, as determined by a PCA plot of the SNPs called.

After the data were downloaded from NCBI, adapter sequence removal and quality score filtering of FASTQ reads was performed using Trimmomatic v0.36 (Bolger et al. 2014) using the parameters “LEADING:28 TRAILING:28 SLIDINGWINDOW:8:28 SLIDINGWINDOW:1:10 MINLEN:65 TOPHRED33”. Trimmed reads from each polyploid sample were mapped to the 26 chromosomes of the *G. hirsutum* reference genome (Saski et al. 2017), and reads from each diploid sample were mapped to each subgenome separately to avoid competitive mapping of the diploid reads against a tetraploid reference genome. Reads from the three outgroup species were separately mapped to both subgenomes to ensure that reads were not filtered out for mapping to multiple parts of the genome. All mapping was done using bwa-mem v0.7.17 (Li and Durbin 2009) and only uniquely mapping paired reads (-F 260 flag) that were mapped in their proper orientation (-f 2 flag) were retained using Samtools v1.9 (Li et al. 2009) before the files were sorted and converted to bam files. Using the Sentieon (Kendig et al. 2019) SNP Calling program, gVCF files were generated, and joint genotyping was performed using the GVCFtyper algorithm (see Github repository for full scripts). SNP filtering was performed using GATK v4.0.4.0 using the filter expression “QD < 2.0 ∥ FS > 60.0 ∥ MQ < 40.0 ∥ SOR > 4.0 ∥ MQRankSum < −12.5 ∥ ReadPosRankSum < −8.0”. For each species (excluding the outgroup species, and treating *G. stephensii,* and *G. ekmanianum* as a single species), we nullified any SNP call in which all individuals were heterozygous to remove any collapsed genomic region in the reference genome or paralogous regions that were not present in the reference genome. We treated *G. stephensii* and *G. ekmanianum* as a single species because we only sampled two individuals of *G. stephensii*, so removing any sites in with both individuals were heterozygous errantly removed real SNPs that were not due to paralogy mapping issues. All scripts for generating and filtering SNP calls are located on our GitHub repository (https://github.com/conJUSTover/Deleterious-Mutations-Polyploidy).

### Identification of Homoeologs

We used the pSONIC pipeline (Conover et al. 2021) to identify syntenically conserved homoeologs in the *G. hirsutum* reference genome, and kept only homoeologous pairs that were less than 5% different in their total annotated CDS length. To remove homoeologous pairs that may have experienced homoeologous exchange events (though there is scant evidence for this (Salmon et al. 2010; Flagel et al. 2012; Chen et al. 2020)), we removed any pair in which the proportion of the reads from the two progenitor diploid genomes (termed At and Dt in the allopolyploid, the “t” indicating “tetraploid”) did not meet the expected 2:2 ratio. Average read depth of CDS regions was determined by bedtools2 v.2.27.1 (Quinlan and Hall 2010). Briefly, for a single homoeologous pair, we calculated the average read depth of the At homoeolog divided by the sum of the average read depth of both homoeologs and removed any homoeologous pair in which this fraction was less than 37.5 or greater than 62.5. We expect any HEs that result in a 0:4 At:Dt copy number to contain 0% At reads/total reads; HEs that result in 1:3 At:Dt copy number should have a 25% At reads/total reads; HEs that result in a 3:1 At:Dt copy number should have a 75% At reads/total reads; HEs that result in a 4:0 At:Dt copy number should have a 100% At reads/total reads; and no HE (i.e. 2:2 At:Dt copy number) would result in a 50% At reads/total reads. We used the midpoints between the “No HE” and the 1:3 and 3:1 copy numbers as cutoff points. This filtering resulted in 8,884 homoeologous pairs (17,768 genes) being analyzed further.

Non-reciprocal homoeologous exchanges (i.e. homoeologous gene conversion) could also bias the estimates of the genetic load in a way that is not related to new mutation following polyploidization or speciation. To control for positions in these non-HE homoeologs that may be influenced by gene conversion, we linked SNP positions between homoeologs in the following way. We first performed pairwise alignments of the CDS sequences using MACSEv2 (Ranwez et al. 2011; Ranwez et al. 2018), which aligns CDS sequences in accordance with their translated amino acid sequences, but allows for the possibility of frameshift mutations. We then used the aligned CDS sequences to identify where indels were present, and found the corresponding genomic positions for every nucleotide in the alignment, inserting gaps where indels occurred.

We then extracted the genomic positions for each SNP position as well as the genomic position for its aligned nucleotide. We retained only those homoeologous SNP positions in which both positions had a confidently called ancestral allele (described above) and in which the ancestral allele matched between the two homoeologs. Importantly, for homoeologs that were encoded in opposite orientations in the reference genome (i.e. one homoeolog was encoded on the forward strand of the reference genome, and the other homoeolog was encoded on the reverse complement), we ensured that the inferred ancestral states for the two SNP positions included both nucleotides of a purine/pyrimidine pair (e.g. the ancestral state for homoeologous SNP was “A” while the ancestral state of the other homoeologous SNP was “T”). We also removed any pair of homoeologous SNPs in which more than 2 alleles were present (while similarly treating homoeologous pairs encoded in opposite directions as described in the previous sentence).

In total, we only used those SNP sites that: (A) did not link to an indel in its homoeologous pair, (B) were biallelic and had consistently inferred ancestral states in the two subgenomes, (C) the derived allele was found in only one of the two subgenomes or their respective diploid progenitors, and (D) the derived allele was fixed in a diploid and segregating in its respective subgenome (or vice-versa).

### Quantifying Deleterious Mutations

We used GERP++ (Davydov et al. 2010) to identify regions of the genome that are evolutionarily conserved, using whole genome alignments from 11 genomes spanning the Eudicots (Supplementary Table 2). Species were chosen if they contained chromosome-level assemblies publicly available on Phytozome or NCBI, and if all documented whole genome duplication events in each species’ evolutionary history is also shared by *Gossypium* (e.g. the *Arabidopsis thaliana* genome was not chosen because it has experienced at least one independent WGD event since its divergence from *Gossypium*). Genomes were aligned to the *G. hirsutum* reference genome using the LASTZ/MULTIZ approach used by the UCSC genome browser. Briefly, genomes were masked using Repeatmasker using a custom repeat library enriched with *Gossypium* TEs (Grover et al. 2017). Each query genome was aligned to each of the *G. hirsutum* reference chromosomes separately. These alignments were chained together using axtChain, and the best alignment was found using ChainNet. These alignment files were converted into fasta files using the roast program from the MULTIZ package.

Using these genome alignments, we used the gerp++ package (Davydov et al. 2010) to calculate GERP scores for every position in the genome. First, we used 4-fold degenerate sites in all genomes to calculate a neutral-rate evolutionary tree, which was calculated using RAxML (Stamatakis 2014). We then used the gerp++ package to estimate the GERP score at every position in the genome, but importantly, we excluded the *G. hirsutum* reference genome from the alignment to avoid biasing sites in the reference genome that may be deleterious. Because the gerp++ program ignores gaps in the reference genome, we used custom R scripts to enter dummy variables in the gapped regions of the GERP score so the number of GERP scores equaled the total number of nucleotide positions in each chromosome. Scripts for each step above are available on Github (link here). To calculate the genetic load across linked sites, we used the GERP load (i.e. the sum of the derived allele frequency times the GERP score for each SNP site) as described in (Wang et al. 2017) and (Rodgers-Melnick et al. 2015). All scripts for generating the multiple sequence alignments and GERP scores can found in our GitHub repository (https://github.com/conJUSTover/Deleterious-Mutations-Polyploidy)

Secondly, we used the BAD_Mutations (Kono et al. 2016; Kono et al. 2018) pipeline to perform LRT tests on conserved amino acid substitutions sites. Nonsynonymous substitutions were identified using SNPEff (Cingolani et al. 2012) and statistical significance was determined using a Bonferroni correction with 967,155 missense mutations to correct for multiple testing. Every step of the BAD_Mutations pipeline was performed using the dev branch of the github repository (accessed July 13, 2020). Species included in the calculation of deleterious mutations are included in Supplementary Table 3, with the notable absence of Gossypium raimondii since it was sampled as part of this project.

We used the GERP load (sum of the allele frequencies * GERP score) (Wang et al. 2017) and the BAD_Mutations load (sum of the allele frequencies of all statistically significant deleterious mutations) as a summary of the genetic load present in each genome at different phylogenetic depths. The BAD_Mutations load may be interpreted as the average number of deleterious alleles expected in each individual of a population, but it does not differentiate between severity of deleteriousness (as does GERP load). We also used GERP to classify SNPs into mildly deleterious (0<GERP≤2), moderately deleterious (2<GERP≤4), and strongly deleterious (4<GERP≤6). Scripts for generating the whole-genome alignments for GERP are located on our GitHub repository (https://github.com/conJUSTover/Deleterious-Mutations-Polyploidy).

### Rate of Deleterious Mutations Along the Phylogeny of Gossypium

To determine if there was a bias in the rate of deleterious mutation accumulation between the two subgenomes, we used homoeologous SNPs in which the derived allele showed a parsimony-informative position between the two subgenomes of allopolyploids and the two diploid progenitors (identified by the green bars in Supplemental Figure 1).

### Genetic Diversity

For each species, *π* for the 17,768 high quality gene CDS sequences (8,884 homoeologous pairs) was calculated on a site-wise basis using vcftools (Danecek et al. 2011). To find the total PI across all genes, we summed the total sitewise pi values and divided by the total length of the concatenated CDS sequences, removing any positions which did not have a null SNP call in the VCF file.

## Acknowledgements

The authors thank the Iowa State University ResearchIT Unit for computational support. The authors also thank Guanjing Hu for her assistance in designing Figure 1, and Matthew Hufford for his helpful comments and discussions on the manuscript. This work was supported by funding from the National Science Foundation-Plant Genome Research Program (awarded to JFW) and Cotton Inc. (awarded to JFW and JLC).

---

**Supplementary Figure 1:**
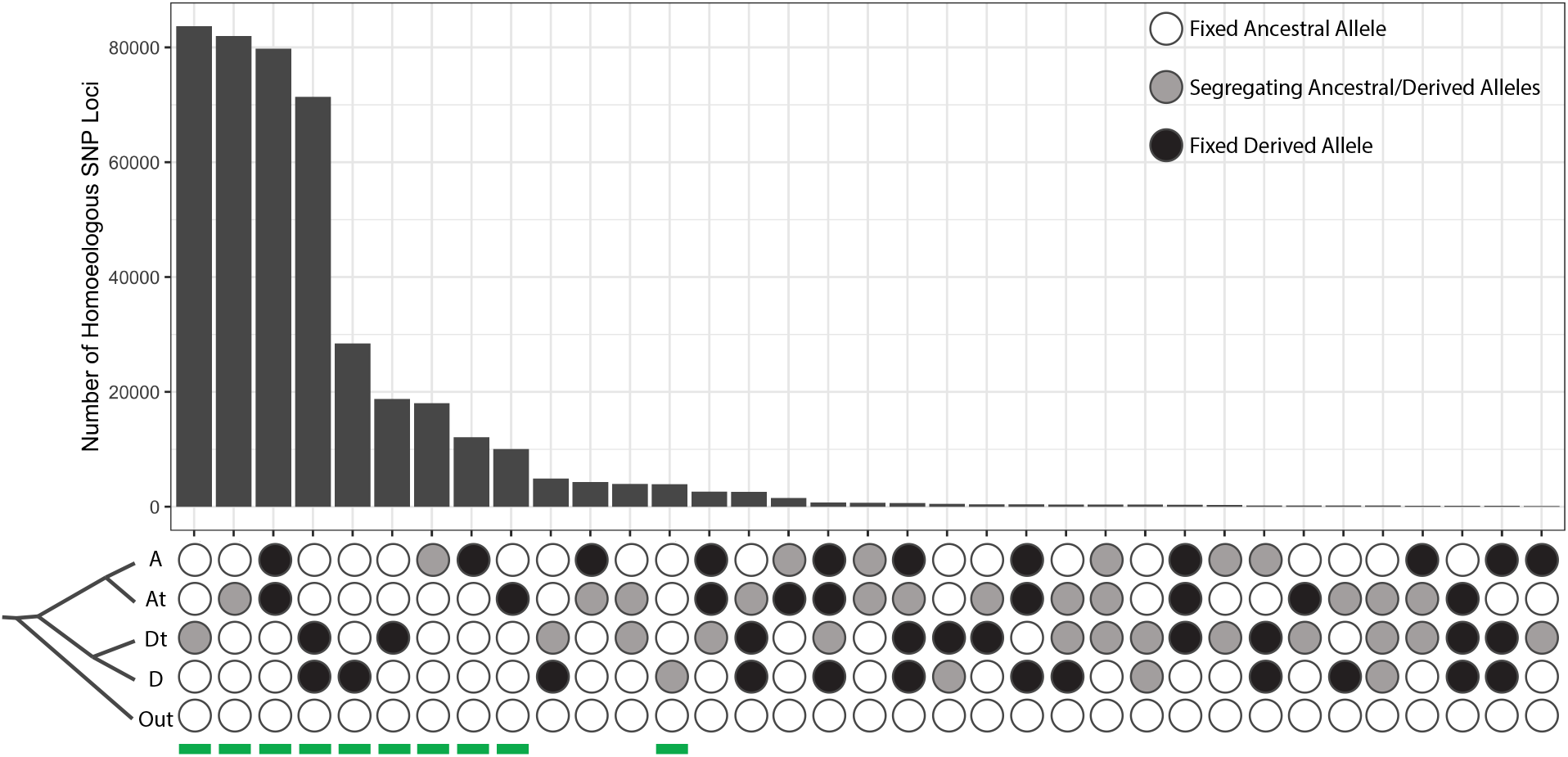
UpSet Plot of Derived Homoeologous SNPs Among 8,884 Syntenic Homoeologous Gene Pairs. To identify SNPs that may have potentially arisen from causes other than simple nucleotide substitutions (e.g., sequencing error, gene conversion), we plotted the frequency of polarized (ancestral vs derived) SNPs across the four major clades of *Gossypium* allopolyploid genomes (A diploid, At subgenome, Dt subgenome, D diploid). Bottom of the UpSet plot shows the phylogenetic positions of these 4 groups, as well as the ancestral state used for polarization. For simplicity, we collapsed all polyploids into a single group, but split them by subgenome (e.g. the At row indicates the At subgenome in all 6 allopolyploids in this analysis). White bubbles indicate that only ancestral alleles were identified in that species or subgenome; black bubbles denoteSNP sites where only derived alleles were identified; grey bubbles represent SNP sites where both ancestral and derived alleles were identified. Only the top 35 SNP groups are shown. Groups with a green line underneath indicate SNP patterns that can be explained by a single mutational event with no homoplasy (e.g. from incomplete lineage sorting or recurrent mutation), and were retained for subsequent analyses involving the 8,884 homoeologous gene pairs.

**Supplementary Figure 2:**
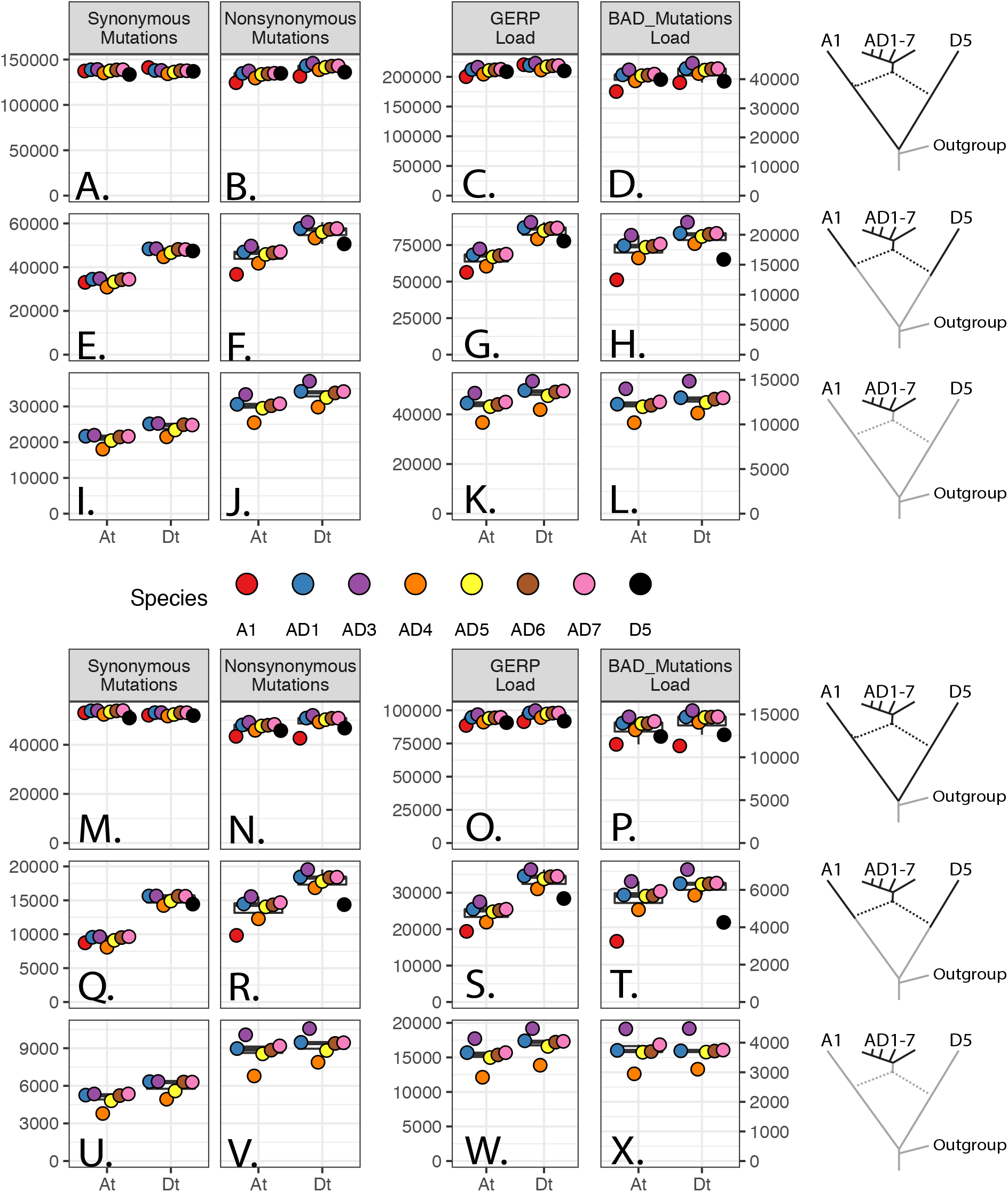
Genome-Wide Derived Mutations and Deleterious Loads at Three Phylogenetic Depths. Number of derived synonymous, nonsynonymous, and deleterious mutations in the CDS regions of 8,884 pairs of homoeologs (17,768 genes in total) in eight cotton species at three phylogenetic depths (indicated by bold branches of phylogeny at right). For all panels, the ancestral state of each SNP was determined using three Australian cottons as an outgroup (see Methods). The deepest phylogenetic depth **(ABCD)** includes all derived mutations that originated since the divergence of the A and D diploid progenitors; the middle row **(EFGH)** shows SNPs that are variable within each subgenome and its associated progenitor diploid species; and the bottom row **(IJKL)** shows SNPs that originated post-polyploidy and are variable within the polyploids. **(AEI)** Synonymous mutations. **(BFJ)** Nonsynonymous mutations. The y-axis for both synonymous and nonsynonymous is shown at left, and represents the sum of the derived allele frequencies, interpreted as the average number of derived SNPs in that category in each species. **(CGK)** GERP Load of each species, calculated as the sum of (derived allele frequency * GERP Score) for all SNP positions with GERP > 0. **(DHL)** Number of deleterious mutations in each species, calculated by BAD_Mutations with bonferroni corrected significance (see Methods). Y-axis represents the sum of the derived allele frequencies, and indicates the average number of deleterious mutations in each species at a given phylogenetic depth. **Note:** for **(EFGH)**, comparisons between subgenomes cannot be made because the D5 diploid is more distantly related to the D subgenome than the A1 diploid is related to the A subgenome. Therefore, we would expect a larger number of derived mutations in D than A simply due to evolutionary history rather than to polyploidization *per se*. The panels above the figure legend are identical to those presented in Figure 2. The panels below the figure legend **(M-X)** follow the same order as **(A-L)**, but represent the genome-wide totals without any filtering based on homoeologs or potential sites that are due to gene less, mapping biases, or homoeologous gene conversion and is provided to demonstrate that our filtering criteria did not have a noticeable impact on the patterns of SNPs that we observed, and that homoeologous interactions have a minimal effect on patterns of evolution following allopolyploidy in *Gossypium*.

**Supplementary Figure 3:**
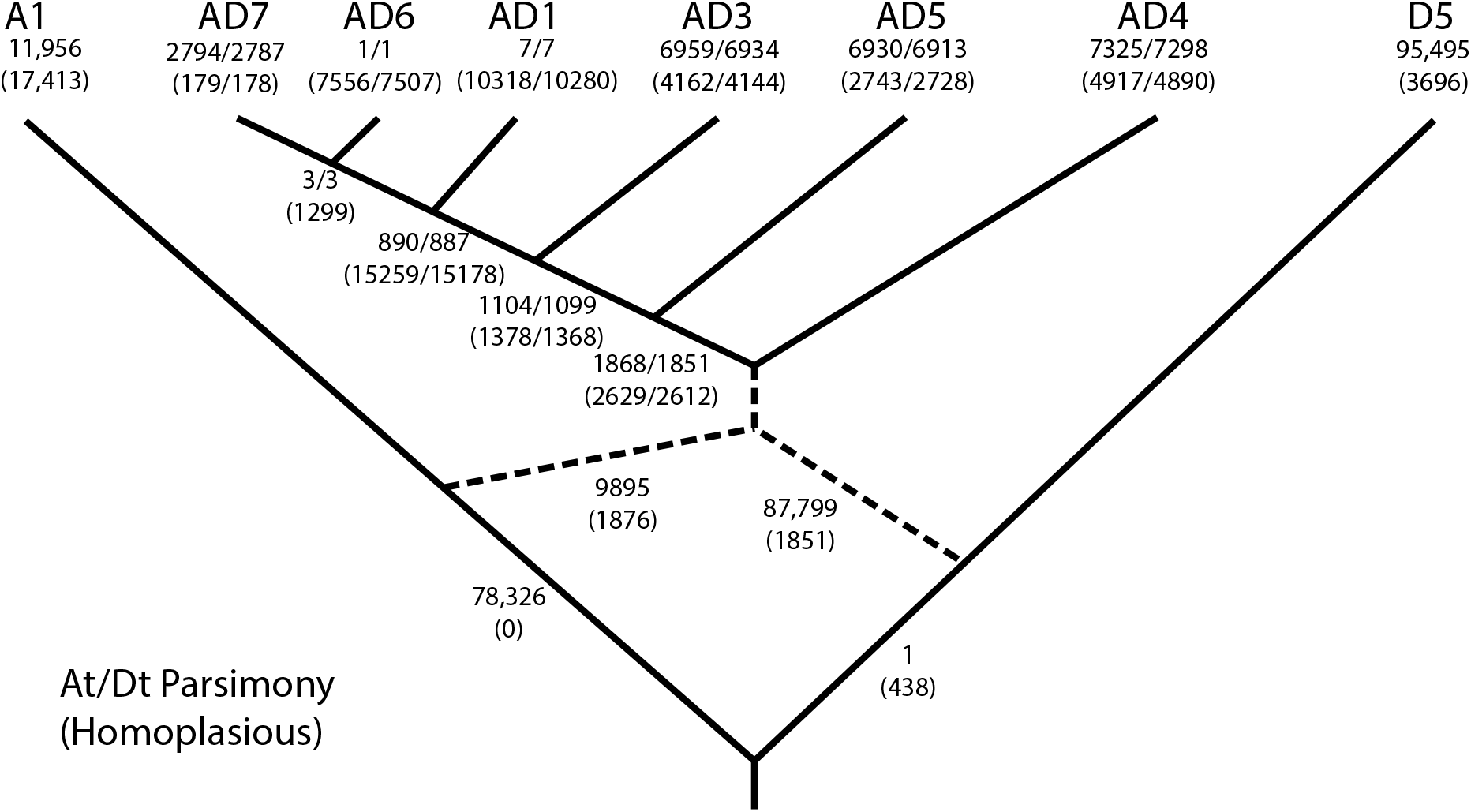
Phylogenetic Positions of Derived Deleterious SNPs. For SNPs that passed the filtering from Supplementary Figure 1, we placed the origin of the SNP on the phylogenetic tree using parsimony. Numbers in the format of “X/Y” indicate the number of SNPs found in the “At/Dt” subgenome. Numbers above the parentheses indicate SNPs that are unequivocally placed on the tree in either the At or Dt subgenome. Numbers in parentheses indicate SNPs that are homoplasious, and the position of the number represents the phylogenetic position of the most recent common ancestor of all species that contain at least one derived SNP. Numbers in the parentheses at the tips of the tree indicate SNPs that are segregating within that species but are not found in any other species. Note: the high amount of homoplasious SNPs at the base of the AD1, AD6, and AD7 clade is most likely caused by recent hybridization or introgression of AD1 into AD6, as also indicated in Supplementary Figure 5.

**Supplementary Figure 4:**
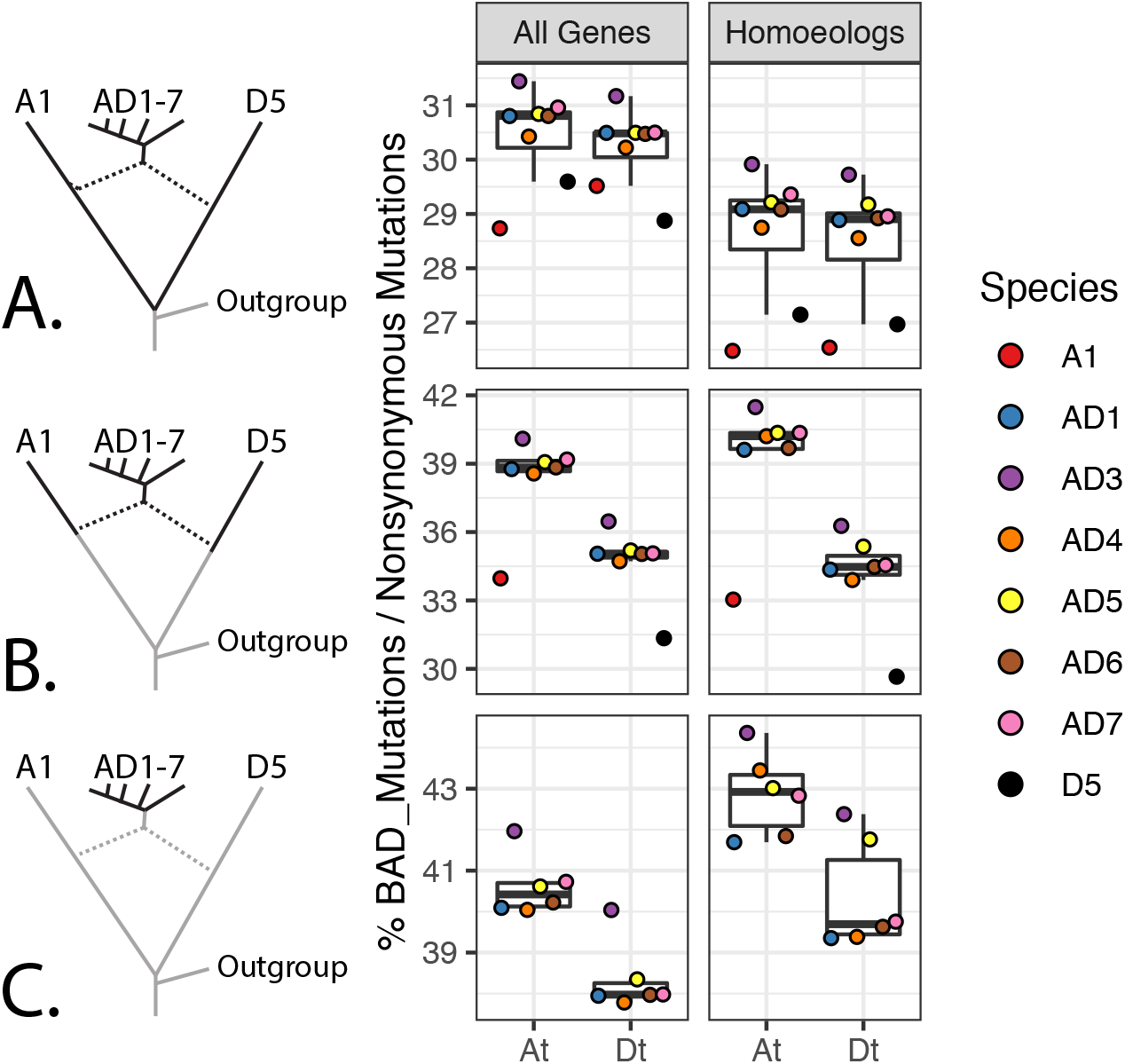
Genome-Wide Proportions of All Nonsynonymous Mutations That Are Deleterious. Rows **A, B**, and **C** summarize SNPs segregating within the entire clade, within each subgenome and its respective progenitor diploid, and within each subgenome, as indicated by the bolded branches along the phylogeny at left. **(A)** Proportion of all nonsynonymous SNPs that are deleterious genome-wide within each subgenome. **(B)** Proportion of nonsynonymous SNPs that are deleterious within 8,884 homoeologous pairs (17,768 total genes) that are syntenically conserved between the two subgenomes of *G. hirsutum* (see Methods for filtering criteria). Note: Similar to Figure 2, comparisons between subgenomes in row **B** reflect differing phylogenetic distances, not asymmetries between the subgenomes and/or their diploid progenitors.

**Supplementary Figure 5:**
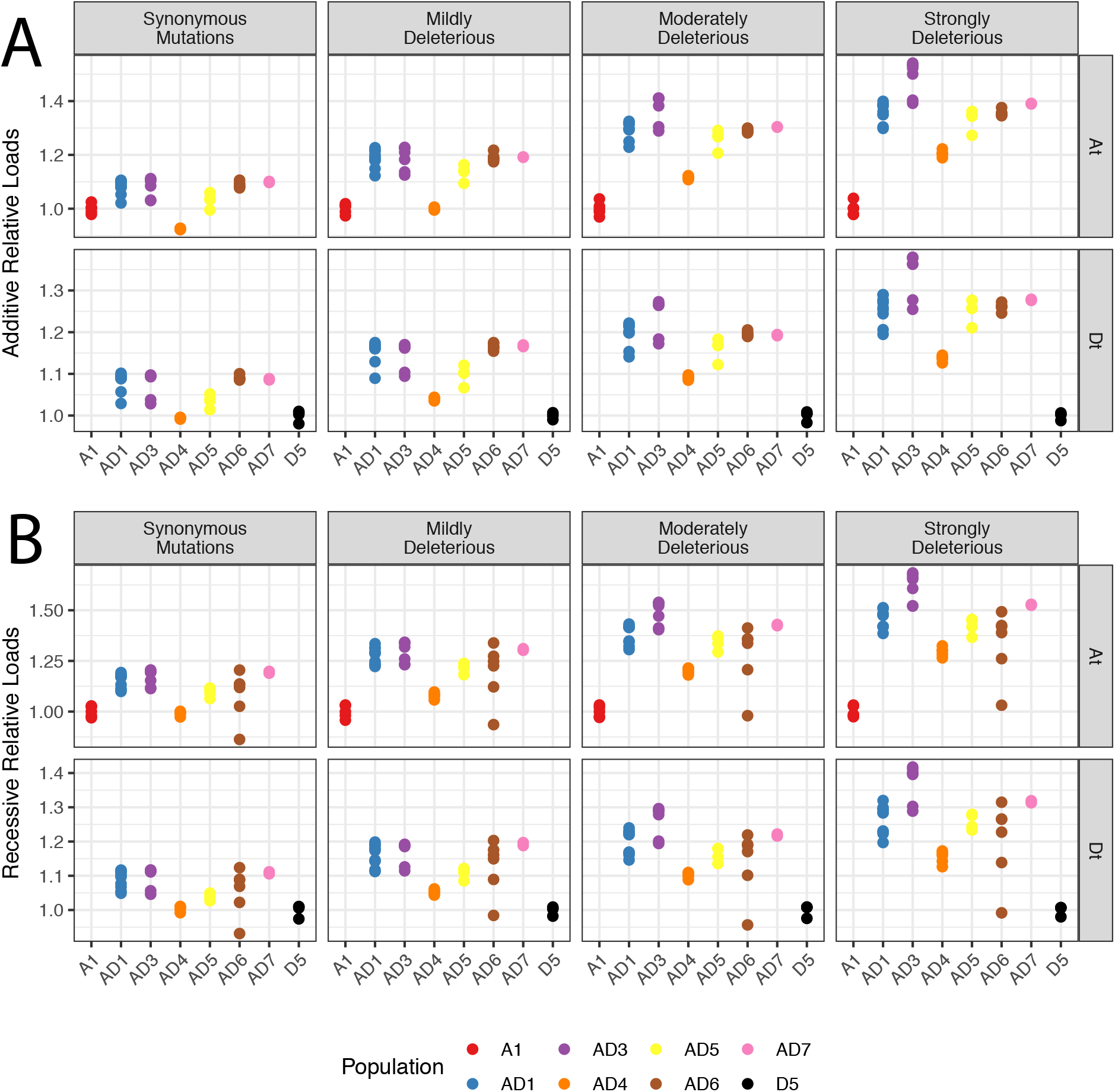
Additive and Recessive Models of Deleterious Mutation Accumulation. Relative load of synonymous sites and varying GERP categories from an **(A)** additive model (i.e. counting all SNPs) and **(B)** recessive model (i.e. counting all homozygous SNPs in a homozygous state). Each point represents an individual, and the placement of each point represents the relative increase or decrease in the number of SNPs relative to the average of the number of SNPs in the diploid (A1 for At, D5 for Dt). Note: The high variance in the recessive load for AD6 reflects a high number of sites that are heterozygous. This is mostly likely due to recent hybridization or introgression from AD1, which is also indicated by a high amount of incomplete lineage sorting between AD1, AD6, and AD7 in Supplementary Figure 3.

